# Estimating epidemic incidence and prevalence from genomic data

**DOI:** 10.1101/142570

**Authors:** Timothy G. Vaughan, Gabriel E. Leventhal, David A. Rasmussen, Alexei J. Drummond, David Welch, Tanja Stadler

## Abstract

Modern phylodynamic methods interpret an inferred phylogenetic tree as a partial transmission chain providing information about the dynamic process of transmission and removal (where removal may be due to recovery, death or behaviour change). Birth-death and coalescent processes have been introduced to model the stochastic dynamics of epidemic spread under common epidemiological models such as the SIS and SIR models, and are successfully used to infer phylogenetic trees together with transmission (birth) and removal (death) rates. These methods either integrate analytically over past incidence and prevalence to infer rate parameters, and thus cannot explicitly infer past incidence or prevalence, or allow such inference only in the coalescent limit of large population size. Here we introduce a particle filtering framework to explicitly infer prevalence and incidence trajectories along with phylogenies and epidemiological model parameters from genomic sequences and case count data in a manner consistent with the underlying birth-death model. After demonstrating the accuracy of this method on simulated data, we use it to assess the prevalence through time of the early 2014 Ebola outbreak in Sierra Leone.

## Introduction

A primary goal of infectious disease epidemiology is to understand epidemic dynamics which are most fully described by the prevalence and incidence of cases through time. Yet most epidemics are only partially observed so their dynamics need to be inferred using statistical methods on incomplete data that can come from a wide variety of sources and over a wide range of scales. A key tool for summarising and understanding epidemic dynamics are compartmental models—such as the SIR model [1]—which partition the hosts at any time into compartments (e.g., susceptible, infectious or removed) and describe how the counts in the compartments change. By estimating the parameters of a compartmental model, we can calculate fundamental quantities like the basic reproductive number, *R*_0_, or simulate prevalence and incidence curves to approximate the true epidemic. However, the reliability of these estimated quantities heavily depends on the adequacy of the model used.

In recent years, several statistical methods have been developed for epidemiological inference from genomic data. These methods lie at the intersection of statistical phylogenetics and epidemiology, and exploit the rapid evolution of many pathogens that occurs on the same time-scale as their epidemiological spread. In these cases, pathogens are said to be *measurably evolving* [2] and the use of phylogenetics in this context is termed *phylodynamics* [3].

Early phylodynamic methods used ad hoc methods to infer epidemiological parameters, incidence, and prevalence. The “skyline plot” [4], based on the mathematical relationship between the effective population size and the time between coalescent events in phylogenetic trees [5], was first used to produce non-parametric estimates of HIV prevalence [4]. Later, in the context of Hepatitis C virus, skyline plots were fitted to a parametric epidemiological model to estimate the basic reproduction rate, *R*_0_ [6]. A subsequent approach combined the estimation of the viral phylogeny and the effective viral population size through time into a joint Bayesian method known as the Bayesian skyline plot [7], but this still lacked an explicit model of the epidemiological process. Another variant of the skyline plot based on the birth-death process [8] allowed for piecewise-constant variation in the birth and death rates [9] from which *R*_0_ could be derived. An important limitation of all of these approaches is that they either do not directly integrate epidemiological modeling into the phylogenetic inference method, or they use piece-wise constant approximations to changing incidence and prevalence through time.

There have recently been three approaches to incorporate compartmental models into phylodynamic inference. First, Volz *et al*. [10, 11] showed how to derive prior probability distributions for viral gene trees in the coalescent limit from arbitrary birth-death processes. This method gives a theoretical basis for joint Bayesian inference of epidemic model parameters, prevalence curves and phylogenetic trees. Inference of model parameters and prevalence curves has been performed using this theory [12–14]. The coalescent basis of this method requires epidemic curves to either be deterministic, or stochastic as long as the epidemic events are statistically independent from the events that make up the sampled epidemic transmission tree [12]. Either assumption is justified in the case of large population size (prevalence). But when prevalence is low, the coalescent method is known to lead to biased estimates of the phylogenetic tree and the epidemiological parameters [15,16]. Furthermore, large sample fractions may lead to violation of statistical independence assumption, as in this case the majority of epidemic events are present on the sampled phylogeny.

Second, Kühnert *et al*. [17] used a parametric compartmental model—specifically, a stochastic SIR model—to produce the piecewise-constant rates of the birth-death skyline plot. Like the coalescent methods of Volz *et al*. [10, 11], this enables joint inference of epidemiological parameters, epidemic curves and phylogeny which can be performed using the software package, BDSIR. The stochastic formulation of the epidemiological process does not rest on the assumption of large population sizes but, like the coalescent methods, the tree events and the epidemic events are assumed to be statistically independent.

Third, Leventhal *et al*. [18] presented the first inference approach to employ an approximation-free computation of the phylogenetic tree probability under a stochastic epidemiological model. The method involves a tailored numerical algorithm to integrate the master equations of a stochastic epidemiological process that is conditioned on the phylogenetic tree. While this approach can be extended to full joint inference of epidemic model parameters and the phylogeny, the available implementation assumes a known phylogeny and integrates using differential equations over all possible prevalence curves to infer epidemic model parameters.

In this paper, we introduce a new method that uses the Particle Marginal Metropolis-Hastings algorithm (PMMH) [19] to jointly infer prevalence and incidence curves, phylogenetic trees, and epidemiological parameters under stochastic epidemiological models. Our approach addresses several of the short-comings of previous methods: (i) it accounts for the dependence of epidemic and tree events; (ii) it incorporates stochastic models of epidemic dynamics; (iii) it includes “sampled ancestors”; and, (v) it provides a natural route to the inclusion of additional (non-genetic) incidence data in full joint phylodynamic analyses. The sampled ancestors [20] mentioned in (iii) are samples which appear in the phylogenetic tree as direct ancestors to other samples, meaning a patient transmitted after the time of sampling and one or more patients in the downstream transmission chain were also sampled.

While particle filtering approaches have been previously applied to phylodynamic inference [12, 13, 21, 22], our application is distinct. In the case of Rasmussen et al. [12], this approach has only been used in the diffusion limit where the discrete nature of the compartment occupancies is ignored. This assumption was relaxed in Rasmussen et al. [13], however the tree density was still computed using a coalescent approximation and inference was conditioned on a known genealogy. Similarly, Li et al. [22] employed particle filtering to estimate the effect of non-geometric distributions of secondary infection counts on the estimation of reproductive number under a coalescent assumption. In contrast, our particle filter is used to compute the exact probability of a transmission tree under the full stochastic discrete compartmental model and used within a joint inference framework. This distinction is especially important near the start of epidemics where prevalence is low and diffusion or coalescent limits do not hold [16]. In the case of Smith et al. [21], particle filtering is applied to individual-based epidemic models. Such models offer greater flexibility than the compartment-based models we use here at the expense of greater computational complexity and a correspondingly lower limit on the number of samples that can be realistically analyzed.

Note that in this paper we use *prevalence* to refer to the absolute number of infectious individuals, as this connects concretely to the discrete population models we employ. The proportion (rather than absolute number) of infected individuals can also be easily derived using the methods we describe, as we will demonstrate.

## New Approaches

In this section we derive a flexible and exact inference method for unstructured stochastic compartmental models.

### Stochastic compartmental epidemic models

Compartmental models are the centrepiece of epidemiological modeling. They partition individuals in a population into compartments according to their infection status and describe how they transition between the compartments. For example, in an SIS model individuals are either susceptible (S) or infectious (I). Susceptible individuals move to the infectious compartment upon infection, and infectious individuals move back to the susceptible compartment upon recovery. The SIR is similar to the SIS model, except that infectious individuals do not move back to the susceptible compartment, but are removed (R) meaning that these individuals cannot move back to the infectious compartment. Removal may be due to, for example, recovery with immunity, or death. Let *S*[*t*], *I*[*t*] and *R*[*t*] (or the relevant set for a given model) represent the number of individuals in the respective compartments at time *t*, and define 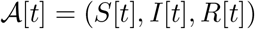 to be the state of the epidemic at time time.

In this paper, we consider unstructured compartmental models: models in which there is only one class of infected individual, i.e. those individuals in the single infectious compartment. This rules out (i) models that include an exposed compartment, often called E, where an individual can be infected but not yet infectious (such as SEIR and SEIS), and (ii) structuring of the infectious compartment via space, age or other factors. The reason for this restriction is that lineages of the transmission tree we discuss below would, under a structured model, require labelling to indicate the compartment each part of the lineage occupies thereby greatly increasing the difficulty of the inference problem.

The overall epidemiological model is defined by the set of compartments, the set of epidemic event types, 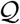, and their corresponding rates, 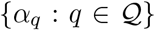. The transitions of individuals between compartments via the epidemic events can be described by a continuous-time Markov process on the state vector 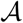 with master equation

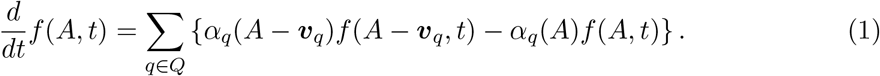

Here 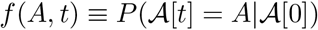 is the probability that the system state 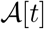 at time *t* has the particular value *A*, *α_q_*(*A*) is the overall rate at which the epidemiological event of type *q* occurs when the epidemic is in state *A*, and *v_q_* is the effect of event type *q* on the state: *A* → *A* + *v_q_*.

This formulation encompasses a broad range of models. For instance, a linear birth-death model consists of just one compartment: 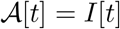, the number of infectious individuals at time *t*. Possible events are infections and removals, so 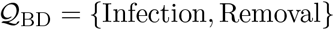. The infection event produces a single new infection as described by *v*_Inf_ = +1, and the overall infection rate is 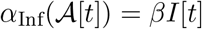. Here, *β* is a constant describing the rate at which infectious individuals produce subsequent infected individuals. Similarly, the removal event removes an individual from the infectious compartment, *v*_Rem_ = −1, at overall removal rate 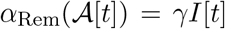. The SIS model, 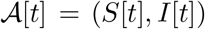, has the same event type set as the linear birth-death process, 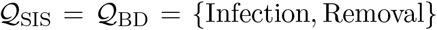, but different rate functions and event effects. An infection has effect vector *v*_Inf_ = (−1, 1) and occurs at rate 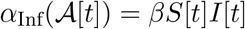, while a removal event has an effect vector *v*_Rem_ = (1, −1) and occurs at rate 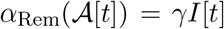. The SIR model, 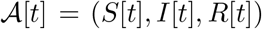, is similar to the SIS model, only with effect vectors *v*_Inf_ = (−1, 1, 0) and *v*_Rem_ = (0, −1, 1). For brevity, we combine the set of constants into a single variable ***η, η***_BD_ = ***η***_SIS_ = ***η***_SIR_ = (*β, γ*).

A specific realisation of an epidemic forward in time—an epidemic trajectory—up to some predetermined maximum time *T* can be generated as follows: The epidemic starts at at time *t*_0_ = 0 in state 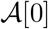. Typically, *I*[0] = 1 for the infectious compartment, but other choices are possible. This initial state is modified by a series of events with types *e*_1_, *e*_2_, …, *e_s_* at times *t*_1_, *t*_2_,…, *t_s_*, where *s* is a random variable indicating the number of events which occurred before *T*. The number of the individuals in each compartment after the *i*-th event has occurred at time *t_i_* is denoted by 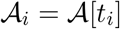. The population trajectory of the epidemic is then just given by 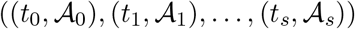. Figure 1a shows an example of the infectious compartment occupancy over time. We can then equivalently expand 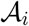 as a sum of effect vectors:

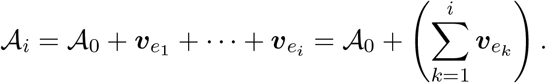

**Figure 1:**
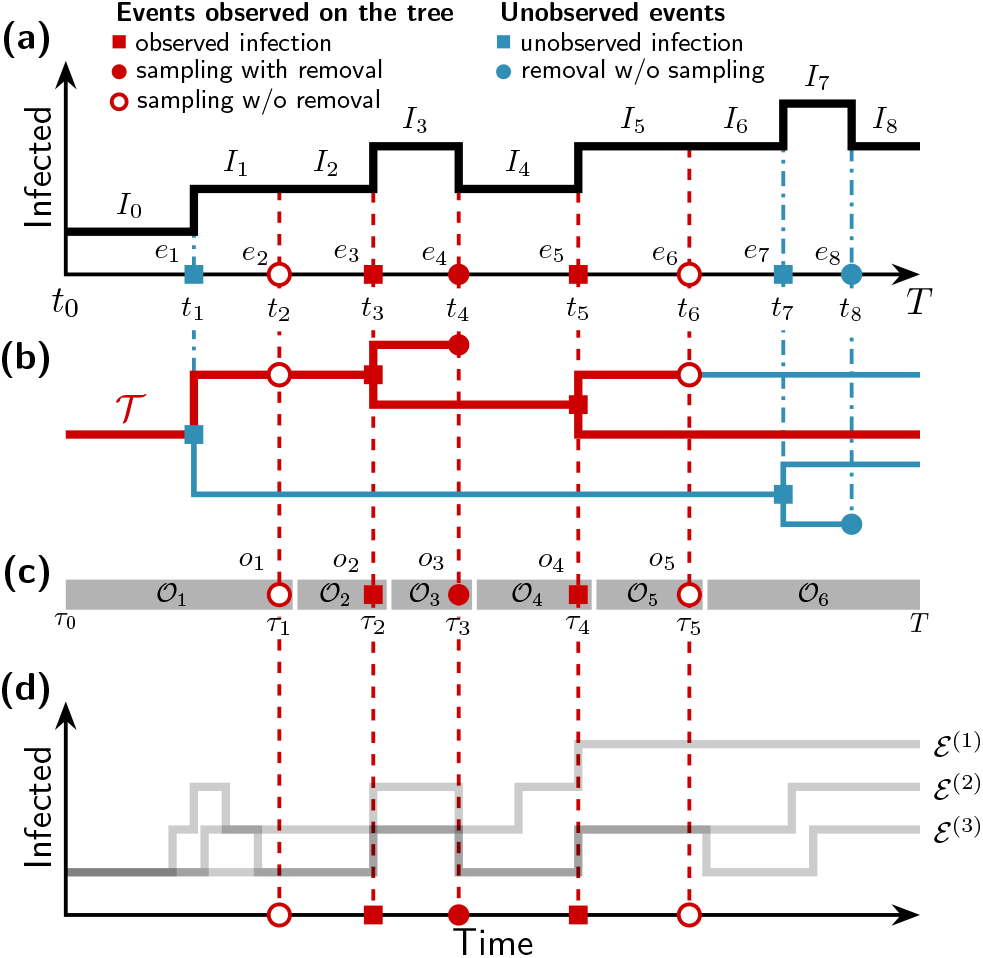
The true epidemiological trajectory can be inferred from the reconstructed phylogeny. **a**. The trajectory ℰ of an epidemic outbreak consists of a sequence of events (infection, sampling, recovery) *e_i_* at times *t_i_* that result in a corresponding sequence of compartment occupancies such as the infectious compartment occupancies *I_i_*. **b**. The full transmission tree contains information on when infections happened and between which lineages (filled squares) and when individuals were removed (filled circles). The sampled transmission tree 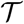 represents a subset of the full tree (red). The rest of the transmission tree is unobserved (blue). **c**. The time ordered observations 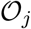 consist of the events o_j_ seen on the tree (infection, sampling w/ removal, sampling w/o removal) at times *τ_j_*, combined with the number of lineages on the sampled tree in the intervals immediately before each of these events. **d**. There is an ensemble of trajectories ℰ^(1)^, ℰ^(2)^,… that are compatible with the sampled transmission tree. Note that the sampled transmission tree contains only a subset of the events represented by the full tree and true trajectory ℰ, and each of these “observed” events must be present in every compatible trajectory.

An epidemiological trajectory ℰ is thus well defined by the initial state, 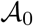, the vector of transition events **e** = (*e*_1_, *e*_2_, …, *e_s_*), and the corresponding event times, **t** = (*t*_1_, *t*_2_,…, *t_s_*),

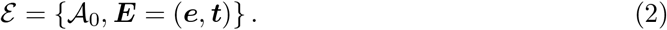

As for any time-homogeneous discrete state continuous time Markov process, the probability density of a particular trajectory is a product of exponentially distributed waiting times between the *s* events with factors representing the probability density of each event. That is,

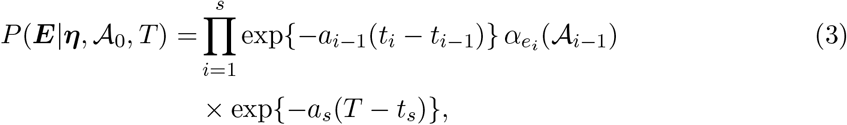

where 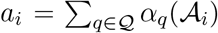 is the sum of the rates of all possible transitions in the interval (*t_i_,t*_*i*+1_). For example, under the SIS model new infections happen at a rate *βS_i_I_i_* and infected individuals are removed at a rate *γI_i_*. By defining 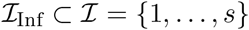 to be the indices of infection events, and 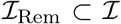 to be the indices of the removal events, we can write the probability density for an SIS trajectory as,

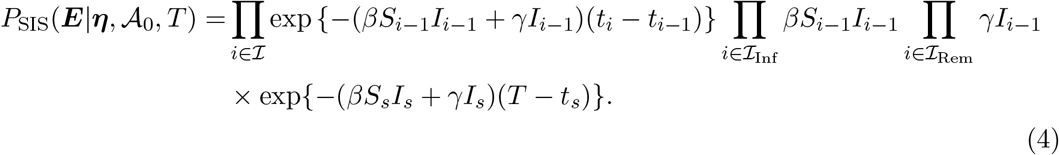

### Modelling the sampling process

Sampling of individuals can be described by expanding 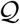 to include two additional event types, sampling with and without removal. While the particular form of the effect vectors depend on the dimension of the compartmental model, their effect remains the same: *v*_SampR_ removes an individual from the infectious class, while *v*_SampNR_ leaves 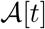 unchanged. We explicitly model sampling by augmenting the stochastic process with sampling events and times, and their corresponding rates: 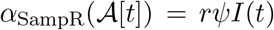 and 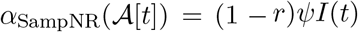, where *ψ* is the per-individual sampling rate parameter and *r* is the probability of removal following sampling. Additionally, any remaining infected individuals at time *T*, i.e. the end of the process, are sampled with probability *ρ*. For convenience, we group all parameters related to sampling together in the vector *σ* = (*ψ,r, ρ*). We then define 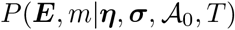 to represent the probability density of this combined process producing a trajectory ***E*** terminated by *m* contemporaneous samples at time *T*. For example, in the case of the SIS model this probability density is

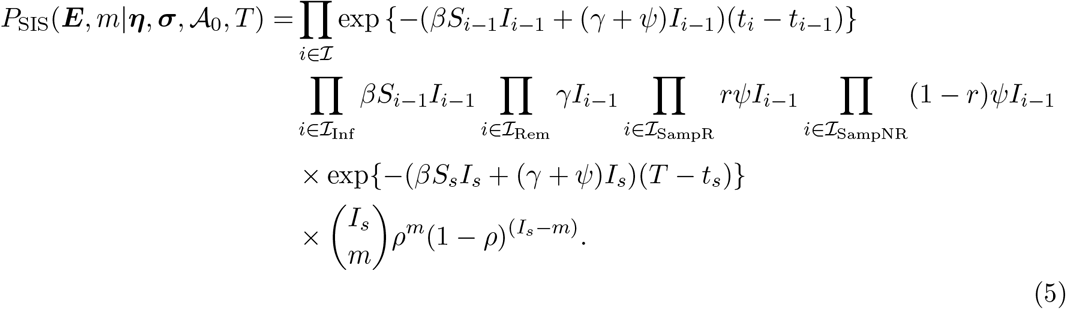

### From epidemiological trajectories to transmission trees

By tracking the identity of who infected whom along an epidemiological trajectory, we obtain the transmission tree of the epidemic (full tree in Figure 1b). All events *e_i_* in the trajectory (Figure 1a) correspond to nodes in the full tree. The number of extant lineages in the full tree *immediately following* the event time *t_i_* is *I_i_*.

The *sampled phylogeny*, 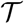, is the subset of the full tree where only the subtree ancestral to sampling events is retained (red subtree in Figure 1b). We use *k_i_* to represent the number of lineages present in the sampled phylogeny *immediately following* time *t_i_*, so *k_i_* ≤ *I_i_*. The number of lineages remaining in the sampled tree at time *T* is *k_s_* = *m*.

Because of its relation to the full tree, each node in the sampled phylogeny must correspond to a compatible event in the trajectory for the probability of the sampled phylogeny given the trajectory 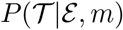 to be non-zero. Furthermore, this distribution is independent of the particular epidemiological model. In particular, conditional on the trajectory, the sampled phylogeny can be considered a result of a discrete-time Markov chain proceeding from the most recent sample to the start of the epidemic process. This can be illustrated by defining 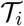 to be the portion of the sampled phylogeny 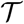 on the interval (*t_i_, t*_*i*+1_], i.e. including the tree node (if any) which corresponds to the event *e*_*i*+1_. We assume that lineages in the tree 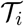 are labelled, such that the correspondence between lineages in 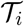 and 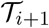 is unambiguous.

For example, an infection event, *e_i_* = Infection, in the trajectory only produces a branching event in the sampled tree when both the infector and the infected correspond to lineages in the sampled phylogeny, so

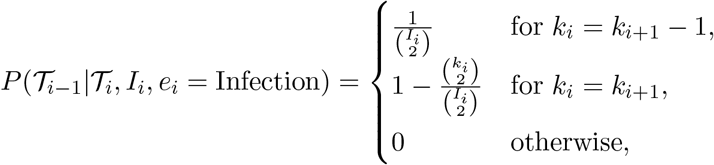

where *I_i_* is the total number of infected individuals (including the newly infected individual) and thus 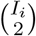 is the total number of pairs of lineages after the infection event, each of which could have been the pair of lineages involved in the event.

Unsampled removal events do not themselves correspond to any nodes in sampled phylogenies, so if *e_i_* = Removal we have

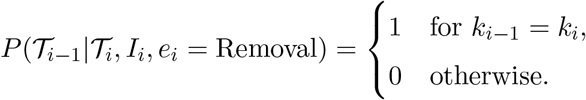

On the other hand, any sampling with removal event corresponds to a leaf node at the time of the event in the sampled phylogeny with probability one:

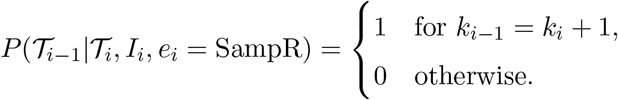

In the case of samples that do not coincide with removal of the sampled lineage, there is ambiguity regarding whether the event is represented by a external leaf node or an internal sampled ancestor node in the sampled phylogeny, as this depends on whether or not any descendants of the sample are subsequently sampled:

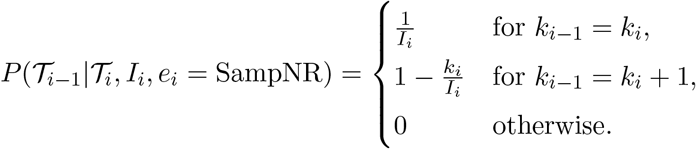

Combining the probabilities above allows us to calculate the full probability of the sampled phylogeny given a complete compatible trajectory as,

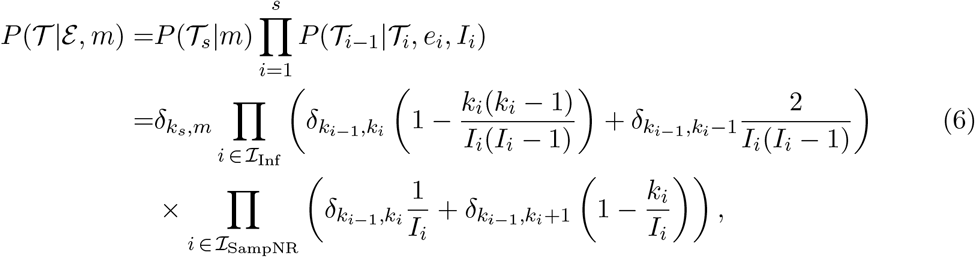

where *δ* is the Kronecker delta, and 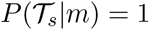 provided *k_s_* = *m*.

### Accounting for unsequenced samples

We now consider the possibility that samples generated by the birth-death-sampling process may be absent from the sampled phylogeny. These samples, which we refer to here as *unsequenced samples*, arise naturally in epidemiological settings where a large number of pathogen samples may be collected at known times but only a subset are subsequently sequenced. Similarly, doctors’ records can provide evidence that individuals were carrying a pathogen at a particular time, but without sequencing there is no information about where exactly the pathogen lineages ancestral to these samples attach to a sample phylogeny.

It is possible to directly include unsequenced samples in the phylogeny but their relationship to the rest of the phylogeny would not be informed by data and they would contribute nothing to the inference of relationships between the sequenced samples while increasing the complexity of the overall inference problem.

Instead, we assume that the set of all sampling event indices 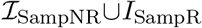 is arbitrarily partitioned into subsets 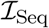 and 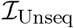 containing indices of sequenced and unsequenced sampling events, respectively. (By allowing this partitioning to be arbitrary, we are choosing not to explicitly model the decision to sequence a given sample, but to instead condition on this decision.) Since this classification then has no effect on the probability density of the stochastic trajectory, we simply exclude the unsequenced sample indices from the final product in the tree probability given by Eq. (6). This gives the following joint probability for the time tree 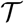 and the unsequenced sample times 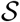:

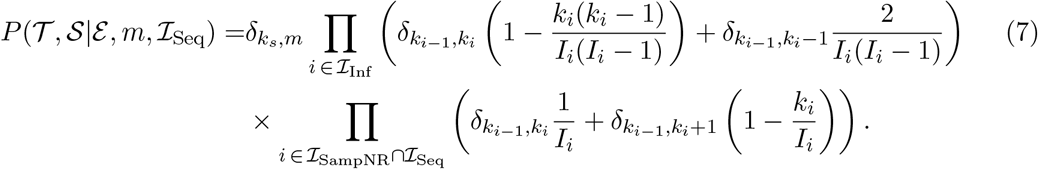

Again, we emphasise that this expression assumes each event in 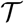 and 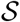 has a corresponding event in the trajectory ℰ and that otherwise the joint probability is zero.

### Bayesian inference

One of our goals is to perform asymptotically exact Bayesian inference of both the prevalence trajectory and the epidemiological parameters using a set of pathogen sample times, a subset for which genetic sequence data are available, collected throughout an epidemic. To this end, for a given pathogen sequence alignment (with a sampling time associated with each sequence) *D* and set of times of unsequenced samples 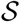, we use Bayes’ rule to express the joint posterior distribution for the model parameters and the epidemic trajectories in terms of the conditional distributions composing the full model:

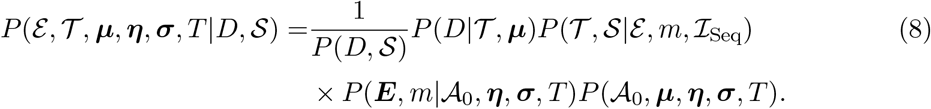

Here 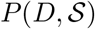 can be treated as a normalisation constant and 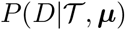 is the probability of *D* evolving down the sampled transmission tree 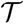 under a substitution model parameterised by ***μ***, also known as the *phylogenetic likelihood*. 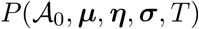 represents the joint prior probability distribution for the model parameters.

Several approaches to characterising this posterior for particular models already exist in the literature, all of which involve using Markov chain Monte Carlo (MCMC) to sample (or maximum likelihood to optimise) a marginalized and/or approximate form of Eq. (8). For instance, Stadler et al. [23] analytically marginalise over the trajectory sub-space in the case of the linear birth-death model and use MCMC to sample from 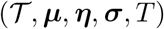. Similarly, Leventhal et al. [18] express the marginalization of Eq. (8) over trajectories for the nonlinear stochastic SIS model as the solution to a master equation which is then integrated numerically with parameter inferences being drawn by applying MCMC or maximum likelihood.

Kühnert et al. [17] provide an approximation to the posterior for discretised trajectories under the stochastic SIR model and use MCMC to sample 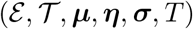. Volz et al. [10] and Volz [11] present an approximation to this posterior under the assumption that the relative amplitude of the stochastic noise in ℰ is negligible and that 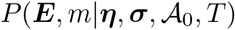 therefore collapses to a point mass centred on the approximate deterministic solution of the model.

In contrast to these methods, we use the particle marginal Metropolis-Hastings (PMMH) algorithm [19]. This has previously been applied in a phylodynamic context by Rasmussen, Ratmann & Koelle [12] and Rasmussen, Volz & Koelle [13] using a coalescent approximation to the distribution of sampled phylogenies, but not to sample directly from the exact phylodynamic posterior as we do in the algorithm described below.

### Particle filtering algorithm

We employ the PMMH algorithm described by [19]. In the form presented here, it involves using a bootstrap particle filter to simulate trajectories ℰ conditional on both a sampled transmission tree 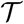 and the times of unsequenced samples 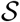.

We call the union of the sampled phylogeny 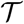 and the temporally distributed unsequenced samples 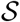 the *observed process*, 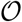, and use *o_j_* to represent the *j*^th^ observation (either a node of the sampled phylogeny or an unsequenced sample) when ordered according to the observation times *τ_j_*, as illustrated in Figure 1c. The final (*N*^th^) observation represents the contemporaneous sampling of *m* lineages in the sampled phylogeny, although it is possible for *m* to be zero.

We divide the time into intervals between observations. The first of these intervals begins at time *τ*_0_ = *t*_0_ = 0, while the last ends at time *T*. We denote the portion of the observed process within interval *j* using 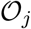, which is understood to include both the number of tree lineages extant within the interval and the observation *o_j_* at end of the interval. Similarly, we divide the full trajectory ℰ into corresponding partial trajectories ℰ_*j*_ which contain only the trajectory events within each observation interval, and define 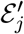 to be the partial trajectory excluding the event *e_j_* corresponding to the observation *o_j_*.

The algorithm involves simulating an ensemble of *M* trajectories or “particles” in each of the *N* intervals between *τ*_0_ and *τ_N_* = *T*. The initial condition for each particle is sampled from the ensemble of finishing states of particles simulated in the previous interval, weighted according to the probability of the observation event that divides the intervals.

The algorithm is as follows:

1. Set the interval index *j* ← 1 and define 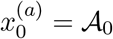 to be the starting state of particle *a*.
2. For each *a* ∈ [1 … *M*] use Gillespie’s stochastic simulation algorithm [24, 25] or its asymptotically exact equivalent [26] to sample a partial trajectory 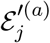 from the distribution

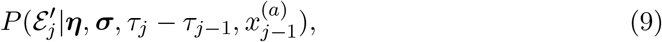

which is a solution to the master equation given in Eq. (1) conditioned on the initial state *x*^(*a*)^ and the interval time *τ_j_* − *τ*_*j*−1_.
3. Each sampled partial trajectory 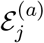 which is defined as the union of 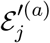 and the event corresponding to the observation *o_j_*, is assigned a weight

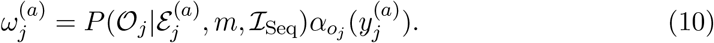 The probability on the right-hand side is given by Eq. (7) but restricted to include only the epidemic events within the interval and the observation event *o_j_*. The factor 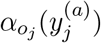 is the transition rate for the epidemic event corresponding to *o_j_* given the final state of 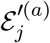, denoted here 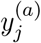. (This factor ensures that the particle trajectories are constrained to be consistent with the observation event *o_j_*, as inconsistent trajectories will be assigned a weight of zero.)
4. The mean of weights 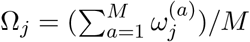 is recorded, and a new set of *M* trajectory states 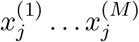 is sampled with replacement from the weighted distribution of the final states of the partial trajectories ℰ_*j*_.
5. If *j* < *N*, set *j* ← *j* + 1 and go to step 2.
6. Compute the product 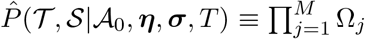 which is, as highlighted below, an estimate of the marginal density 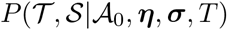, with the marginalization being over the epidemiological trajectories. Also, sample a single final partial trajectory 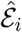 from the final distribution of weighted partial trajectories and follow the sequence of events back through the observation intervals until *t* = 0, yielding a single sampled full trajectory 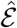.

It can be shown [27] that the value of 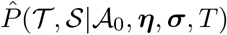 is an unbiased and consistent estimate of the marginal probability density for the sampled phylogeny and unsequenced samples 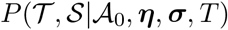. (This probability density is sometimes called the *phylodynamic likelihood*, and below we simply write “likelihood”, although the implicit classification of 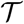 as “data” should not be understood to mean that phylogenies are physically observed.) As shown by [19], this implies that by using this estimate in place of the terms 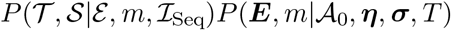 in the posterior given by Eq. (8), and using the resulting expression as the target distribution for a Markov chain Monte Carlo algorithm, we obtain an algorithm for sampling from the joint posterior marginalized over the epidemic trajectories. Furthermore, by recording the sampled trajectories 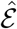 generated by the particle filter alongside the parameter values and sampled phylogenies visited by the MCMC procedure, the algorithm generates samples from the full (unmarginalised) joint posterior.

The use of particle filtering to condition the epidemic trajectories on the tree is potentially confusing, due to the (backward-time) correlations between the observations that make up the sampled phylogeny. Despite these correlations, the PMMH algorithm remains applicable since the joint probability of the observations and hidden state, 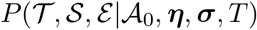, can be expressed in precisely the same form as the weighted sequence of conditional probabilities generated by a standard hidden Markov model. This is shown in the supplementary text, along with a simple demonstration that the resulting algorithm does indeed produce samples from the required marginal density of the observations given the phylodynamic model parameters.

## Results

### Implementation and validation

We have implemented the algorithm described above as a BEAST 2 [28] package. This allows the algorithm to be used in conjunction with standard phylogenetic models such as those describing the nucleotide substitution process as well as existing algorithms for performing the MCMC sampling of the phylogenetic tree space. The package is released under the GNU General Public License and instructions for installing and using it can be found, along with source code, at http://tgvaughan.github.io/EpiInf.

All of the BEAST 2 input files necessary to reproduce the results described in this section, together with instructions on how to use them, may be downloaded from http://github.com/tgvaughan/ParticleFilterResults.

### Direct likelihood comparison

We validated our algorithm and its implementation by comparing the likelihoods generated by the particle filter with those computed analytically under the linear birth-death model [8] and numerically under the nonlinear stochastic SIS model [18]. These comparisons were performed for a variety of parameter combinations and in all cases yielded perfect agreement (Figure 2).

**Figure 2:**
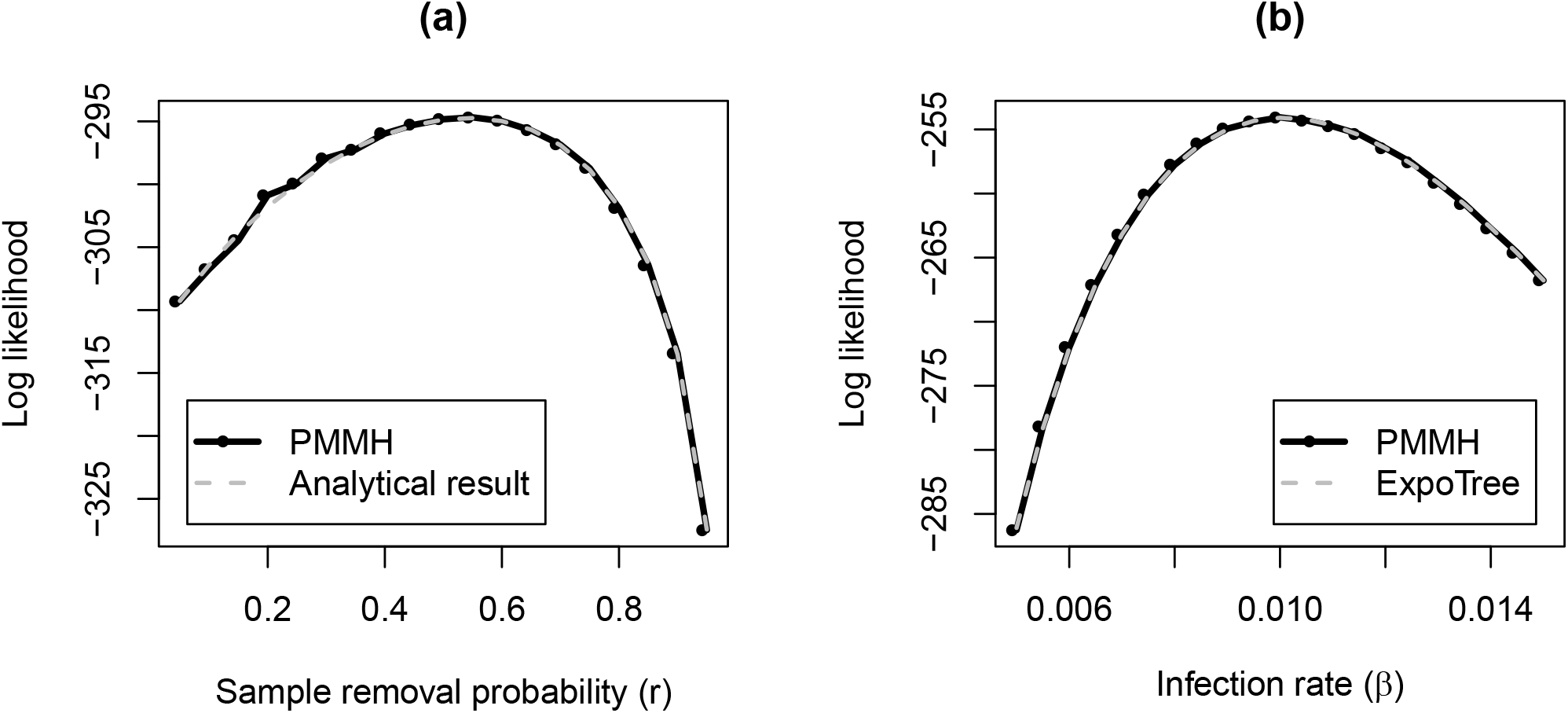
Comparison between values of the phylodynamic likelihoods computed using the PMMH algorithm with those calculated using other approaches: (a) likelihood of *r* under the linear birth-death model from PMMH compared with the analytical result [8] and (b) likelihood of *β* under the stochastic SIS model from PMMH compared with a numerical result from ExpoTree [18].

### Comparison of tree-based and incidence-based sampling

The joint tree and sample time prior defined in Eq. (7) has the property that marginalising over the time tree yields a quantity which is independent of which samples are sequenced and which samples are not. In other words, if the sequence data from the sampled individuals provide no information about the phylogenetic tree then the only information we have are the sample times: our estimates of the epidemiological model parameters should therefore not depend on which samples were sequenced. This suggests the following test for the consistency of the joint posterior:

1. Fix a set of sampling times.
2. Assign a fraction *f* of these times to be associated with tree leaves (i.e. play the role of “sequenced” sample times),
3. Sample from the joint posterior defined in Eq. (8) without sequence data (i.e. setting 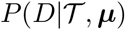 to a constant).

Provided the unsequenced sampling times are being handled consistently by the sampler, the posteriors for model parameters should be identical regardless of *f*.

We performed this test using a set of 83 sample times simulated using a birth-death-sampling process and using these times, via the procedure above, to produce the posterior for the birth rate parameter *β* as a function of *f*. The lack of variation in this posterior as with respect to *f*, shown in figure 3, is strong evidence that our treatment of unsequenced samples is indeed consistent with our treatment of sequenced samples.

**Figure 3:**
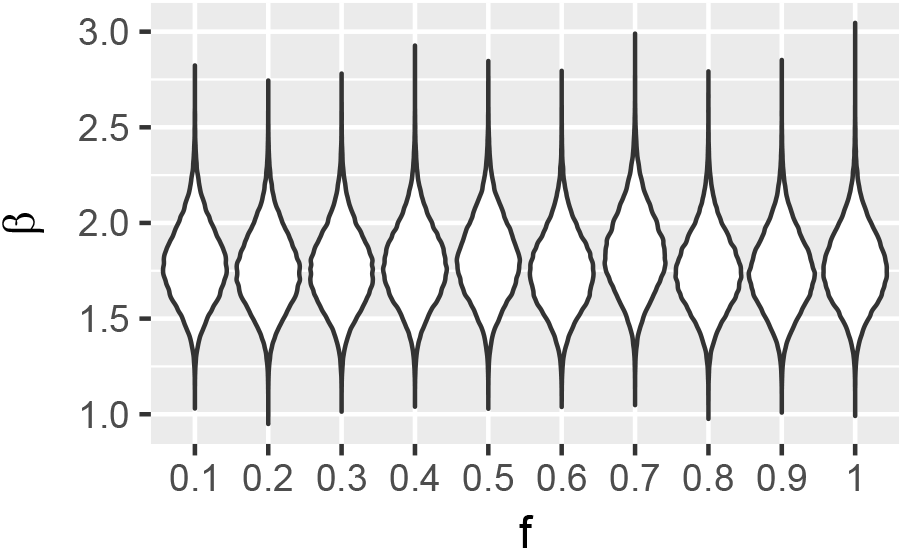
Marginal posteriors for the infection rate as a function of the fraction *f* of samples regarded as “sequenced” when no data besides the sampling times is available. The invariance of this distribution with respect to *f* shows that the treatment of unsequenced samples is consistent with the treatment of sequenced samples.

### Inference from simulated data

In order to assess the capability of the sampler to recover prevalence trajectories, we simulated trajectories under each of the three models supported by our implementation: linear birth-death (*β* = 1.2, *γ* = 0.1, *ψ*/(*ψ* + *γ*) = 0.5, *T* = 7.0), stochastic SIS (*β* = 0.02, *γ* = 1.0, *ψ*/(*ψ* + *γ*) = 0.1, *T* = 5, *S*_0_ = 199) and stochastic SIR (*β* = 0.2, *γ* = 1.0, *ψ*/(*ψ* + *γ*) = 0.1, *T* = 5, *S*_0_ = 199). In all cases we fixed the removal probability *r* = 1, the present-day sampling probability *ρ* = 0 and set *I*_0_ = 1. Sampled transmission trees were then simulated from each of these trajectories, which were in turn used to simulate 2 kb genetic sequence alignments under a simple Jukes-Cantor model with a substitution rate of 5 × 10^−3^ per site per unit time. For each of these three alignments, we then used our algorithm to sample from the joint posterior for the transmission tree, epidemic trajectory and the model parameters *β, γ, T* and (in the case of SIS and SIR) *S*_0_. (The remaining parameters *ψ, r*, and *ψ* were fixed to the truth.) For the continuous parameters we employed improper priors *P*(*β*) = 1/*β, P*(*γ*) = 1/*γ* and *P*(*T*) = 1/*T*. For the discrete *S*_0_ parameter we used *P*(*S*_0_) =Unif(0, 300).

Figure 4 illustrates the agreement between the posterior prevalence distributions obtained from each of these analyses (red lines) and the true prevalence curves (black lines). Also shown is the distribution of prevalence curves generated directly from the posterior samples of the model parameters (blue lines). Prior to our PMMH algorithm, the blue lines were the best estimates obtained for prevalence under compartmental models (unless coalescent approximations were appropriate in the particular application). As these blue trajectories are not explicitly conditioned on the corresponding sampled transmission trees however, there is a significantly greater variance in their distribution.

**Figure 4:**
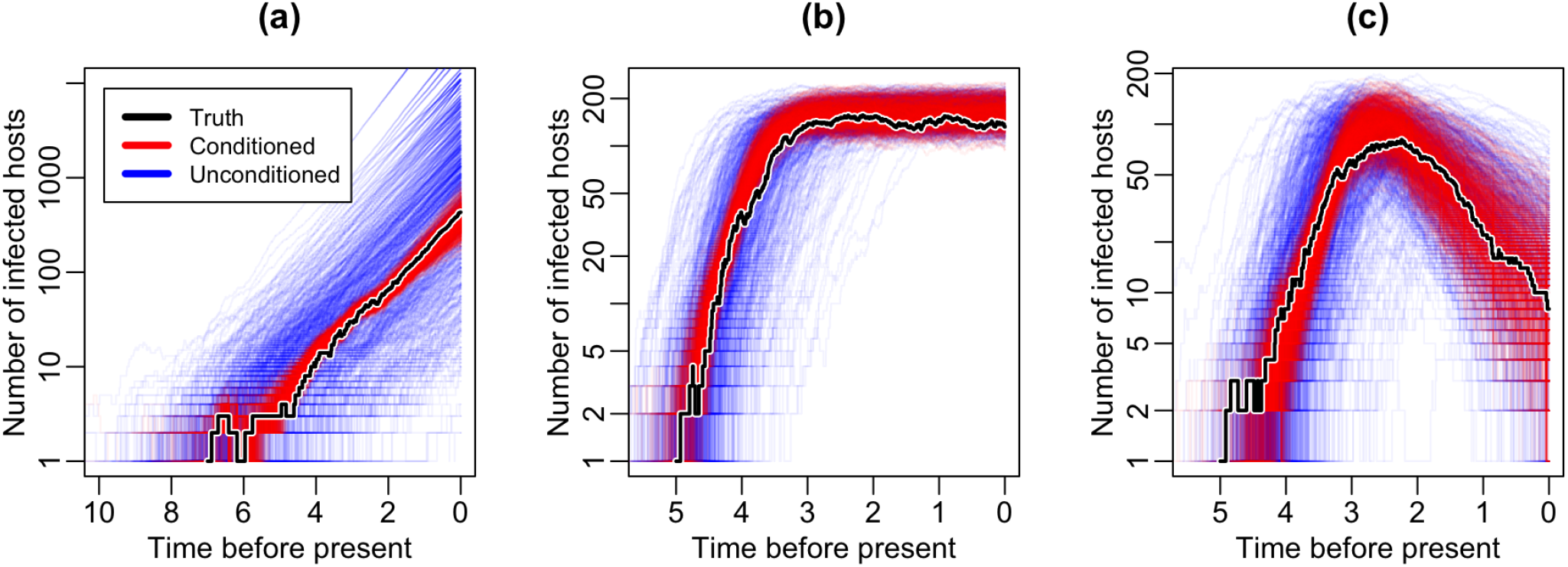
Inference of prevalence dynamics from sequence data simulated under (a) linear birth-death, (b) stochastic SIS and (c) stochastic SIR model. Samples from the posterior of the prevalence trajectory are shown in red, while the black line represents the truth. The blue lines are prevalence trajectories simulated from the posterior samples of the compartmental model parameters.

### Quantitative validation of trajectory inference

While agreement between simulated and subsequently inferred trajectories is encouraging, we use a well-calibrated approach [29] for a more robust quantitative validation of the inference algorithm. The steps of this approach are as follows.

1. Under each model (linear birth-death, SIS and SIR) and a chosen set of parameters (Table 1) we simulate 200 trajectories and sampled trees.
2. A random DNA sequence is simulated down each sampled tree, resulting in a unique simulated sequence alignment.
3. For each simulated sequence alignment, infer the corresponding trajectory conditional on the true model parameters using our inference algorithm.
4. We compute the proportion of analyses for which the true prevalence at a particular time falls within the 100α% highest posterior density (HPD) interval of the sampled posterior distribution for the prevalence at this time. This is repeated for a range of times and α values.

**Table 1:** Fixed parameter values used for well-calibrated trajectory inference validation.

| Model | $\beta$ | $\gamma$ | $S_0$ | $\psi$ | $r$ | $\rho$ | $T$ |
| --- | --- | --- | --- | --- | --- | --- | --- |
| Linear birth-death | 0.5 | 0.1 | — | 0.25 | 0.0 | 0.0 | 10.0 |
| SIS | 0.02 | 1.0 | 199 | 0.1 | 0.0 | 0.0 | 5.0 |
| SIR | 0.02 | 1.0 | 199 | 0.1 | 0.0 | 0.0 | 5.0 |

Figure 5 shows, for each model, the perfectly linear relationship between α and the proportion of analyses for which the 100*α*% HPD includes the truth. This relationship provides strong evidence that our implementation of the algorithm correctly samples from the true distribution of epidemiological trajectories.

**Figure 5:**
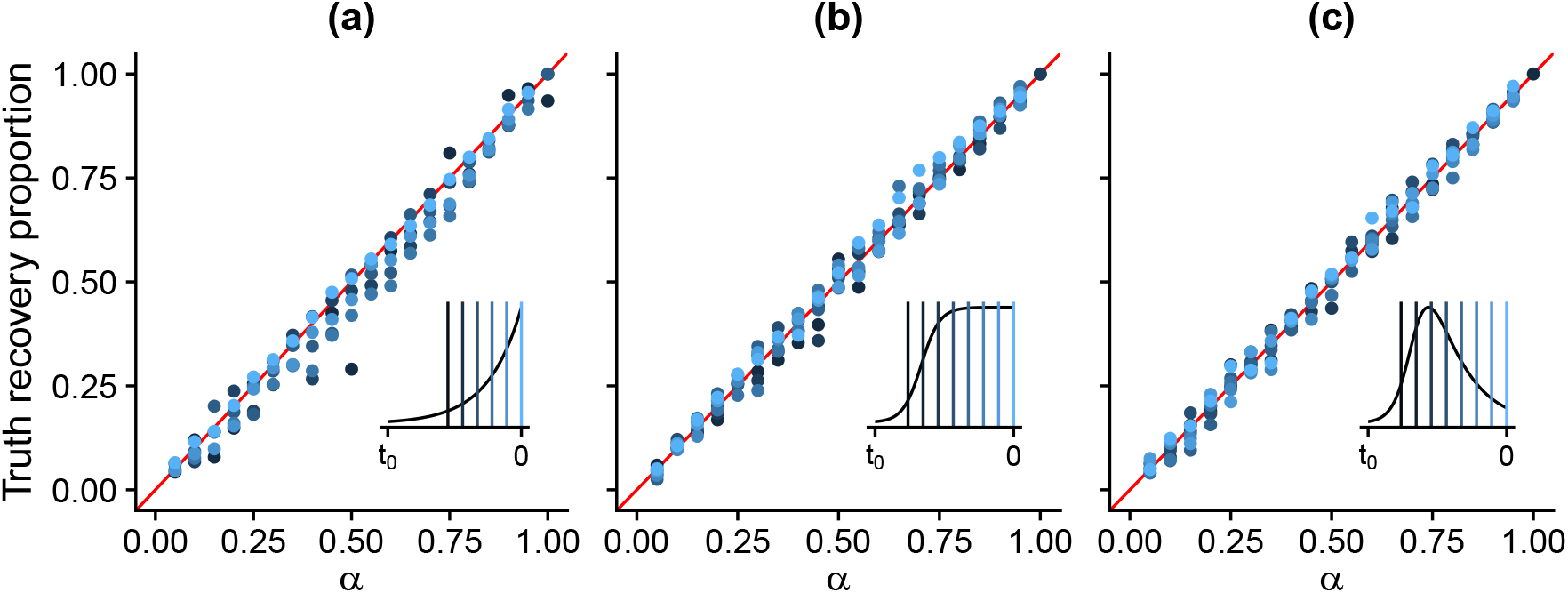
Proportion of simulated data analyses which included the true prevalence in their 100*α*% highest posterior density (HPD) intervals, for alignments simulated under each of the (a) linear birth-death, (b) SIS and (c) SIR models. Colours represent the distinct times at which the coverage fractions were computed, and the insets indicate where these times fall in relation to the approximate deterministic prevalence curves. The linear relationship between the relative inclusion frequencies and α indicates that the PMMH algorithm is correctly sampling from the posterior prevalence distribution under each of these models.

### Inference of Ebola prevalence in Sierra Leone

In order to demonstrate the applicability of our method, we analyzed 101 full Ebola virus (EBOV) genomes collected from the Kailahun district in eastern Sierra Leone during the 2014 west-African epidemic [30–33], as curated and aligned by Dudas et al. [34]. These sequences were analyzed jointly with the temporal distribution of unsequenced Kailahun cases [35]. To assess the degree to which the inclusion of unsequenced data affected the inferred trajectory distributions, we conducted a separate analysis based solely on sequence data collected during the first four weeks. Later sequences were excluded from the latter analysis to avoid introducing bias due to the sequencing fraction being skewed toward earlier weeks (Figure 6f).

**Figure 6:**
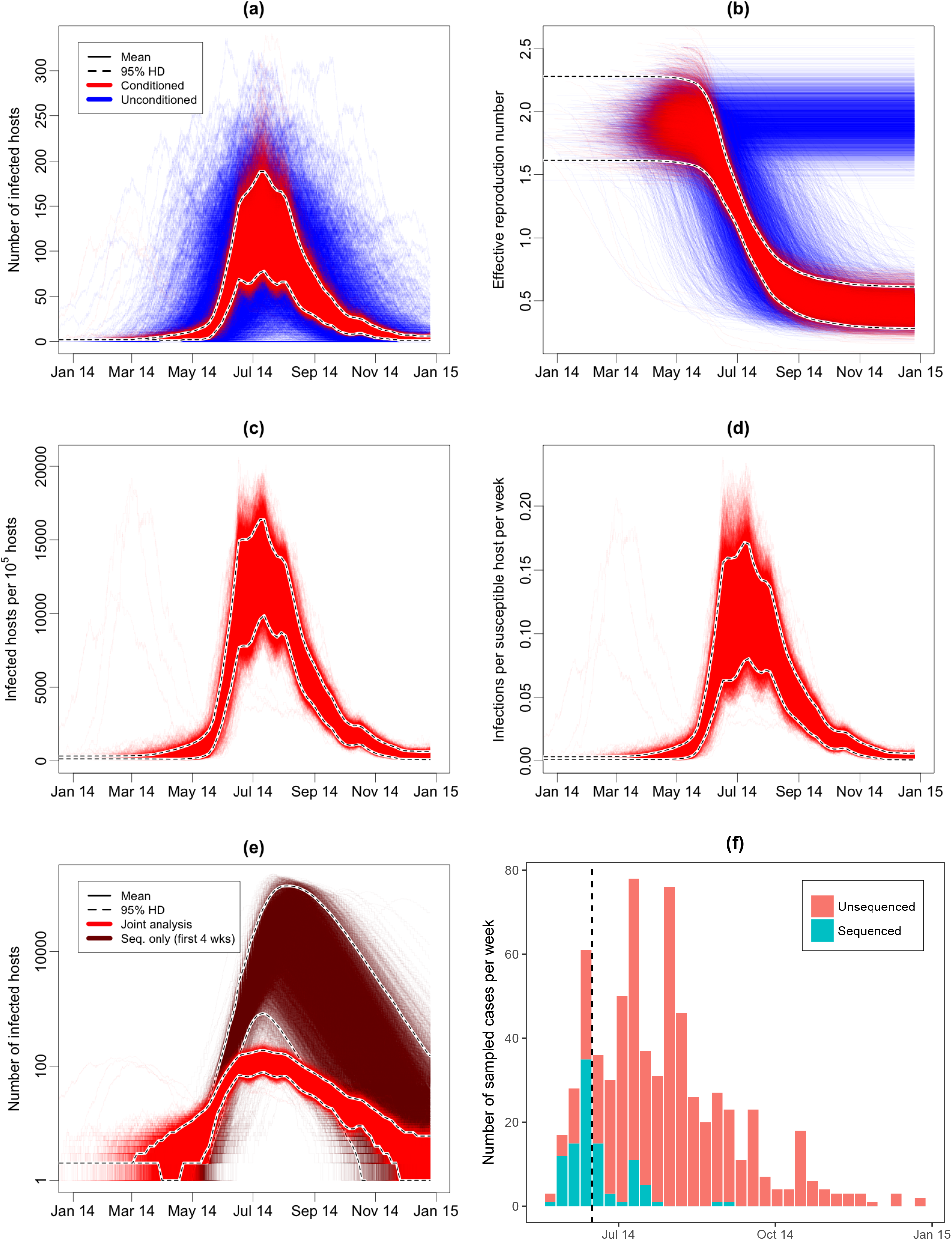
(**a**),(**b**) Jointly inferred posterior distributions (red) and unconditioned simulated distributions (blue) for (a) infected host count and (b) effective reproduction number during the Kailahun EVD outbreak. (**c**) Posterior distribution of infected host count per 10^5^ hosts (prevalence). (**d**) Expected number of of new EVD infections per susceptible host per week (incidence). (**e**) Comparison of inferred number of infected hosts using all data (red curves) and only the first four weeks of sequence data (brown curves). (**f**) Temporal distribution of EBOV cases used in the full analysis, both sequenced (turquoise) and unsequenced (orange). The vertical dashed line in (f) indicates the end of the 4-week period of sequence data used to infer the brown trajectories in (e).

We assumed a standard neutral model of sequence evolution allowing for distinct transition/transversion rates and non-equilibrium base frequencies [36], together with Gamma-distributed rate heterogeneity among sites [37]. We further assumed a strict clock rate whose value was jointly estimated using an informative prior derived from a recent meta-analysis [38].

We assumed a stochastic SIR epidemiological model in which each sample (whether sequenced or unsequenced) is assumed to be generated by a linear sampling process with fixed rate *ψ* between the times of the most recent and earliest samples. Importantly, while the temporal distribution of sample collection times is determined by this model, the choice of which samples to sequence is not. We feel that this is a sensible decision, given the non-linear relationship between the sequenced and unsequenced cases.

The total removal rate *γ* was fixed at 25 removals per infectious individual per year, corresponding to an expected infectious period of approximately 15 days. Similarly, the removal probability at sampling *r* was fixed to 0, meaning that sampling was not assumed to affect infectious potential. All other epidemiological parameters were estimated from the data. The complete list of prior distributions used for these analyses is presented in the second column of Table 2.

**Table 2:** Parameter priors distributions used in and 95% highest posterior density intervals derived from our analysis of EBOV genomes sampled from the 2014 EVD outbreak in Kailahun. Note that while *T* is the time difference between the start of the outbreak and the end of the observation period, for a given time of cessation of observation it implies the absolute time of the start of the outbreak, which we provide in the bracketed (2014) dates.

| Parameter | Unit | Prior distribution | Posterior 95% HPD |  |
| --- | --- | --- | --- | --- |
|  |  |  | Lower | Upper |
| $\beta$ | $\text{year}^{-1}$ | $\text{Unif}(0, 1)$ | $2.9 \times 10^{-2}$ | $8.2 \times 10^{-2}$ |
| $S_0$ | — | $\text{Unif}(0, 5 \times 10^5)$ | 576 | 1390 |
| $\psi$ | $\text{year}^{-1}$ | $\text{Unif}(1, 365)$ | 16 | 36 |
| $T$ | year | $\text{Unif}(0, 2)$ | 0.64 (May 5) | 0.83 (Feb 25) |

For the full analysis and the sequence-only analysis, a total of 30 independent MCMC chains were run for 2 × 10^7^ steps each and compared to assess convergence. The initial 10% of each chain was removed to account for burn-in and the remaining samples combined into two long chains (one for each analysis type) from which the final results were derived.

The 95% highest posterior density (HPD) intervals for each of the estimated compartmental model parameters are presented in the right-most columns of Table 2. Interestingly, despite the broad uniform prior, the initial size of the susceptible population is inferred to be very low: on the order of one or two thousand individuals. This is likely due to the effects of population structure, with the fitted value representing the effective magnitude of the susceptible population rather than a demographic count. Additionally, we find that the overall rate of sampling is comparable to the removal rate *γ*, suggesting a relatively high sampling fraction *ψ*/(*ψ* + *γ*) of 39–60% (95%HPD interval) during the period that sampling was taking place, i.e., between the first and the last sample recorded for this region.

The posterior distributions for the absolute number of infectious hosts, *I*(*t*), and effective reproduction number, *R_e_*(*t*) = *βS*(*t*)/*γ*, trajectories are shown as the distributions of red curves in Figures 6a and 6b respectively. The blue curves shown alongside are trajectories simulated under the model using the sampled epidemiological parameter values and not explicitly conditioned on the observed sample data nor inferred transmission trees, hence their broader variance.

Figure 6c shows the posterior for the prevalence in terms of the number of infectious hosts per 10^5^ initially susceptible hosts in the population. Since the SIR model is a constant population size model, this is also just the proportion of the population at any time which is inferred to be infected. Furthermore, since the initial number of susceptible hosts *S*_0_ is jointly estimated, the shape of the estimated curve differs subtly from the absolute infected host count trajectories shown in Figure 6a due to correlations between this shape and the susceptible host count.

Figure 6d shows the posterior for the rate of incidence. Specifically, it shows the inferred rate of new infections per susceptible host per week, with time measured in weeks.

The comparison between analysis of the full data set and the sequence-only analysis (Figure 6e) clearly displays the advantage of including the additional unsequenced case count data. In particular, it is clear that the unsequenced samples (Figure 6f) provide a wealth of information regarding the peak prevalence of the epidemic, a value that is almost completely unresolved in the sequence-only analysis.

## Discussion

The primary strengths of the inference method and associated software presented here are their versatility and exactness. The method jointly samples from the exact posterior of transmission trees, epidemic trajectories and model parameters under compartmental models without needing to make assumptions about the size of the epidemic or the size of the host population. (In contrast, coalescent methods are usually only applicable when population sizes are large.) The current implementation treats SI, SIS and SIR epidemic models but, with only minor modifications, it can be used under any unstructured stochastic compartmental model whose dynamics can be described by Equation (1).

There is also versatility in the type of data the method accepts. Many phylodynamic methods have relied solely on sequence data to inform their models which, while increasingly available, is more costly and scarce than simple case reports. Our method can use cases reports and sequences together. The benefits of including case reports (unsequenced samples) to improving prevalence estimation are clearly shown in the Ebola analysis where the time of the epidemic peak is much more tightly estimated than when the sequences are analysed alone. We also expect that including the case reports could inform the dating of the tree in data sets where the case reports are numerous and only a small number of sequences are available.

The method described here is also applicable to the field of macroevolution where past species richness, i.e., the number of species through time, is a quantity of much interest. Estimates are typically obtained by using sequences from extant species to estimate past speciation and extinction rates which are then used to simulate unconditioned trajectories [39]. As is the case with epidemic trajectories, using our particle filtering tool to fit conditioned trajectories should improve these estimates and make quantification of species richness more precise. Fossil occurrence data has been shown to greatly improve macroevolutionary estimates [40] and are analogous to unsequenced samples, so can be directly incorporated into analyses with our method.

The sampling model we use is relatively simple, with infected samples uniformly taken at a constant rate through the epidemic and the possibility of burst of sampling at the end. This overly simple approach means that data needed to be discarded in the Ebola analysis so as not to bias results. It is feasible to extend the sampling model to more closely reflect how the data is actually collected, for example by modeling changes in collection effort or having multiple bursts of intense sampling and so avoid potential biases introduced by the current model.

The software implementation of the method within the Beast 2 framework means that the default is to estimate the tree along with other parameters, and the full range of standard phylogenetic models can be used to model sequence evolution along the tree.

The flexibility and exactness of the inference relies on simulation to compute Monte Carlo estimates of the probability density of the transmission tree under the model and so comes at a heavy computational cost. While a single density estimate can be made very quickly, when it is run as part of a larger MCMC analysis, estimates must be computed many times for each MCMC step and for hundreds of millions of steps. The number of simulations run at each step is a tunable parameter of PMMH and does not, in theory, alter the accuracy of the result. But there is a trade-off in that reducing the number of stochastic simulations that make up a density estimate increases the variance of the estimate with the result the Markov chain can become “stuck” after an extreme estimate is made, and the mixing rate of the chain is drastically reduced to the point that independent draws from the target posterior are not being produced. There is potential to parallelise the density estimate by running simulations in parallel at each step though with overheads the benefit of this may be marginal. Overall, joint analysis under this method are currently limited to hundreds of sequences.

Another obvious shortcoming of the present algorithm is its inability to handle structure in the population. Structure can originate from spatial segmentation of the host population or from the infection having distinct phases, for example varying degrees of transmissibility or a non-infectious period (such as in the SEIR model). This issue is addressed by Rasmussen, Volz & Koelle [13], although in an approximate way that assumes events in the epidemic trajectory are independent of the events observed in the phylogeny.

Despite these difficulties, we have presented what is to our knowledge the first algorithm capable of exactly inferring epidemiological trajectories jointly with compartmental model parameters using a combination of pathogen sequencing data and case count records. Our method also enables estimates of species richness through time by combining extant species data and fossil occurrences. A focus for future work will be extending this tool to account for population structure and to allow for the analysis of larger data sets in a mathematically exact framework.

## Acknowledgements

The authors thank Louis du Plessis for helpful suggestions. We also thank the New Zealand eScience Infrastructure for access to high-performance computing facilities (http://www.nesi.org.nz). This work was supported by Marsden grant UOA1324 from the Royal Society of New Zealand. TS is supported in part by the European Research Council under the Seventh Framework Programme of the European Commission (PhyPD: grant agreement number 335529). GEL was supported by the Swiss National Science Foundation (162251) and the Human Frontiers Science Program (LT000643/2016-L).

